# An Incrementally Updatable and Scalable System for Large-Scale Sequence Search using LSM Trees

**DOI:** 10.1101/2021.02.05.429839

**Authors:** Fatemeh Almodaresi, Jamshed Khan, Sergey Madaminov, Prashant Pandey, Michael Ferdman, Rob Johnson, Rob Patro

## Abstract

**Motivation:** In the past few years, researchers have proposed numerous indexing schemes for searching large databases of raw sequencing experiments. Most of these proposed indexes are approximate (i.e. with one-sided errors) in order to save space. Recently, researchers have published exact indexes—Mantis, VariMerge, and Bifrost—that can serve as colored de Bruijn graph representations in addition to serving as *k*-mer indexes. This new type of index is promising because it has the potential to support more complex analyses than simple searches. However, in order to be useful as indexes for large and growing repositories of raw sequencing data, they must scale to thousands of experiments and support efficient insertion of new data.

**Results:** In this paper, we show how to build a scalable and updatable exact sequence-search index. Specifically, we extend Mantis using the Bentley-Saxe transformation to support efficient updates. We demonstrate Mantis’s scalability by constructing an index of *≈* 40K samples from SRA by adding samples one at a time to an initial index of 10K samples.

Compared to VariMerge and Bifrost, Mantis is more efficient in terms of index-construction time and memory, query time and memory, and index size. In our benchmarks, VariMerge and Bifrost scaled to only 5K and 80 samples, respectively, while Mantis scaled to more than 39K samples. Queries were over 24*×* faster in Mantis than in Bifrost (VariMerge does not immediately support general search queries we require). Mantis indexes were about 2.5*×* smaller than Bifrost’s indexes and about half as big as VariMerge’s indexes.

**Availability:** The updatable Mantis implementation is available at https://github.com/splatlab/mantis/tree/mergeMSTs.

**Supplementary information:** Supplementary data are available online.

## 1 Introduction

The number of publicly-available raw sequencing experiment data sets has been growing exponentially throughout the past decade NIH (2020). For example, the Sequence Read Archive (SRA) database currently has more than 17 peta-bases of publicly-available sequencing data and is growing with new samples being added to the database on a daily basis NIH (2020); Leinonen *et al*. (2010); Kodama *et al*. (2011).

Such repositories have the potential to be rich resources for exploratory analyses to answer sequence-level biological questions. For instance, given a newly-assembled transcript or a gene, a researcher might wish to find all the prior sequencing experiments that contained that sequence. This question can be classified as a search problem in which a query sequence is searched across thousands of experiments to determine identical or highly-similar sequences. To efficiently answer sequence-search queries, we need to build indexes on repositories of raw sequencing data. Given the overwhelming size and the continuous growth of such repositories, the indexes need to be memory efficient and must support efficient insertion of new sequencing samples.

The problem of sequence search in a large collection of raw sequencing samples was first introduced by Solomon et al. Solomon and Kingsford (2016a), where they suggested the Sequence Bloom Tree (SBT) as a hierarchical data structure for indexing sequencing experiments for “experiment discovery.” Since the publication of this work, many tools, algorithms, and data structures have been proposed to index large collections of short-read sequencing data (Solomon and Kingsford, 2017; Sun *et al*., 2017; Pandey *et al*., 2018; Yu *et al*., 2018; Bradley *et al*., 2019; Bingmann *et al*., 2019; Almodaresi *et al*., 2020; Harris and Medvedev, 2020) improving performance over previous structures or introducing new indexing schemes. These approaches start by breaking the sequences in a given sample into sub-sequences of size *k*, known as *k*-mers, and define the intersection of the query *k*-mer set and each sample’s *k*-mer set as the criterion for determining the likelihood of the presence of the query sequence in the sample.

The majority of the sequence search indexes are approximate with one-sided errors (false positives). That is, all samples that meet the intersection criterion will be returned, but search results may also include some additional “false positives”, i.e. samples that do not meet the intersection criteria. The false positive rate is typically tunable, at the cost of increasing the size of the index.

Two sequence-search indexes, Mantis (Pandey *et al*., 2018; Almodaresi *et al*., 2020) and Bifrost (Holley and Melsted, 2020), are exact, i.e. they are guaranteed to never return any false positives. Due to this exactness property, these indexes can also be viewed as colored de Bruijn graph (CDBG) (Iqbal *et al*., 2012) representations in addition to being sequence-search indexes. The exact indexes represent the same information as other CDBG representations, such as Cortex (Iqbal *et al*., 2012), VARI (Muggli *et al*., 2017), VariMerge (Muggli *et al*., 2019), and Rainbowfish (Almodaresi *et al*., 2017). Such indexes, being fast and exact, can therefore also be used for applications over CDBGs, such as multi-sample assembly, bubble calling, and variation detection.

Many indexes for the sequence search problem are static and do not support adding new samples to the index without rebuilding the whole index. However, rebuilding the index from scratch is costly, especially as the database grows larger. For this reason, static indexes are not well-suited for large and dynamic sequencing repositories.

In this paper, we show how to build a scalable and updatable exact sequence-search index. Our index is an extension of Mantis, which is already one of the fastest (in terms of query and construction speed) and most space-efficient indexes, in addition to being exact. Our solution is based on the generic static-to-dynamic transformation of Bentley and Saxe Bentley and Saxe (1980). This transformation is well understood and widely used in storage systems (Vora, 2011; Chang *et al*., 2008; Lakshman and Malik, 2010). The main requirement for an index such as Mantis to be able to use the Bentley-Saxe transformation is to support efficient merges, i.e., there must be an efficient algorithm that, given two instances of the underlying static data structure, outputs a new instance that represents the union of the two input instances.

We propose an efficient algorithm for merging two Mantis indexes, and tackle several scalability and efficiency obstacles along the way. Our proposed algorithm targets Minimum Spanning Tree-based Mantis (MST-based Mantis) (Almodaresi *et al*., 2020) which uses a compressed color representation that is over an order-of-magnitude smaller than the original Mantis representation. This allows iterative construction of the full index without the need to ever retain the uncompressed color representation for a large number of samples in memory or on disk.

The proposed merging algorithm builds on MST-based Mantis’s construction algorithm, and we address several scalability problems in the construction process in order to create a scalable merging algorithm. One of our main techniques is to use *k*-mer minimizers to break Mantis’s *k*-mer map into smaller partitions, similar to the scheme used in BCALM2 (Chikhi *et al*., 2016). This enables us to avoid loading a large, monolithic *k*-mer to color-id map in memory during MST construction, merging, and query, reducing the memory bottleneck during large index construction, update and query.

We benchmark the new scalable and mergeable version of Mantis against Bifrost and VariMerge Muggli *et al*. (2017). VariMerge also supports merging, and the VariMerge paper suggests merging can be used to improve scaling of index construction by recursively building indexes on smaller sets of samples and then merging the resulting indexes together. This paper shows that merging is not only useful for improving scalability, but also for supporting updatability.

In our benchmarks, the Mantis merge algorithm outperforms the VariMerge algorithm by 2-4× in all performance metrics measured. Mantis uses a quarter of the memory of the VariMerge merging algorithm and runs in less than half the time. The resulting Mantis index is half as large as the VariMerge index. VariMerge does not expose an interface for *k*-mer point queries, so we cannot compare its point query performance to Mantis’s.

Bifrost does not support merging, so we simply compare Mantis’s construction and query performance against Bifrost’s. Our benchmarks show that Mantis is ≈ 10× faster to construct, requires ≈ 10× less construction memory, results in ≈2.5× smaller indexes, and performs bulk queries ≈74× faster and with ≈100× less query memory than Bifrost.

We also benchmark the query time in Mantis with the partitioned *k*-mer map structure against the previous Mantis representation. We observe similar aggregate query time while obtaining up to a 7× improvement in memory usage for the largest indexes.

Finally, we evaluate our scalable and updatable index, Dynamic Mantis, by measuring performance while adding samples one at a time. We scale the Dynamic Mantis to 39.5K human RNA-seq samples, which is ≈4× larger than previously reported indexes on human samples (10K) and ≈15× larger than the BBB dataset (Solomon and Kingsford, 2016b, 2017; Sun *et al*., 2017; Pandey *et al*., 2018; Almodaresi *et al*., 2020) typically used for evaluation purposes in large-scale sequence search indexes. Our results show the potential of such a data structure to scale to increasingly-larger sets of samples. Combined with a strategy for scaling out across multiple servers, such an approach can eventually target indexing all publicly available human RNA sequence samples.

## 2 Background

We first briefly describe classic and MST-based Mantis data structures. We then explain the Bentley-Saxe construction and describe prior data structures built through this method.

### 2.1 Classic and MST-Based Mantis

Both classic and MST-based Mantis map *k*-mers to the set of samples in which they occur. We call the set of samples in which a *k*-mer occurs the *k*-mer’s “color”. Both versions of Mantis perform this mapping in two steps. In the first step, they map each *k*-mer to a color-class ID. In the second step, each color-class ID is mapped to a color. The first step is performed using a counting quotient filter (CQF), which is a compact hash table for small keys and values Pandey *et al*. (2017b). The CQF also supports efficient enumeration of keys in order of their hash value, which will be useful in our merge algorithm. Both versions of Mantis ensure that *k*-mers that occur in the same set of samples are mapped to the same color-class ID, so the mapping from color-class ID to color is injective. Furthermore, both versions of Mantis assign lower IDs to popular colors, since color IDs are encoded using variable-length counters in the quotient filter.

Classic and MST-based Mantis differ in how they map color-class IDs to colors. In Classic Mantis, color-class IDs are essentially indexes into an array of bit-vectors. Each bit-vector in the array encodes a set of samples. The array of bit-vectors is compressed using RRR (Raman *et al*., 2007), which supports random access to entries in the array. This is similar to the color encoding in Rainbowfish (Almodaresi *et al*., 2017).

MST-based Mantis (Almodaresi *et al*., 2020) further compresses the array of bit-vectors by exploiting similarities between colors. MST-based Mantis organizes the color-class IDs into a tree rooted at the all-zeros vector. It then stores, for each node, the node’s parent ID and the bitwise difference between the node’s bit-vector and its parent’s bit-vector. Thus one can reconstruct the bit-vector for any node in the tree by applying all the bitwise differences along the path from the node to the root. MST-based Mantis constructs the tree by first constructing a weighted graph *C* on all the color-class IDs. For each edge in the DBG connecting *k*-mers *k*_1_ and *k*_2_ it adds an edge to *C*, connecting *k*_1_ and *k*_2_’s color-class IDs. The weight on this edge is the Hamming distance between the IDs’ corresponding bit-vectors. MST-based Mantis then computes an MST for this graph. This approach ensures that the total size of the representation of all the bitwise differences in the MST is minimized. This encoding reduces the size of the color class table by an order of magnitude.

### 2.2 Bentley-Saxe Technique

The Bentley-Saxe transformation is a generic way to transform a static data structure into a dynamic one. The only requirement is an efficient algorithm for merging two instances of the structure into a single one. The transformation maintains *k* = log_*r*_ *N/M* +*O*(1) instances of the static structure, *S*_0_,…,*S*_*k*_, where *N* is the number of items in the structure, *M* is the maximum allowed size of *S*_0_, and *r* is the “fanout”, which can be used to trade-off between insertion and query performance. The value *M* is typically chosen so that *S*_0_ fits in RAM. New items are merged into *S*_0_ until it reaches size *M*. Whenever an instance *S*_*i*_ exceeds size *r*^*i*^*M*, then *S*_*i*_ is merged into *S*_*i*+1_ and *S*_*i*_ is replaced with an empty instance. Queries must search in each *S*_*i*_ in order. Assuming the merging algorithm runs in linear time, then Bentley-Saxes increases the costs of insertions by a factor of *O*(*r*log_*r*_*N/M*) and the cost of queries by a factor of *O*(log_*r*_ *N/M*). Thus small *r* favors insertions and large *r* favors queries. Typical values for *r* in many applications are in the range from 2 to 16.

The Bentley-Saxe construction Bentley and Saxe (1980) has been used extensively in databases and storage systems to build memory-efficient and scalable dynamic systems for indexing large-scale data. For example, log-structured merge trees (LSM trees) (O’Neil *et al*., 1996) are a well known example of using Bentley-Saxe to build scalable databases (Vora, 2011; Chang *et al*., 2008; Lakshman and Malik, 2010).

Bentley-Saxe is particularly useful when data is too large to fit in memory and the merge algorithm performs a linear scan of its input data. In this case, the merge algorithm can be implemented using large sequential I/Os, which make good use of storage bandwidth. This is the case for sorted arrays and static B-trees, the building blocks of LSM trees.

Similar to LSM trees, a cascade filter (Pandey *et al*., 2017b; Bender *et al*., 2012; Pandey *et al*., 2020) is the result of applying the Bentley-Saxe transformation to the (counting or non-counting) quotient filter (Pandey *et al*., 2017c), the data structure used in Mantis to map *k*-mers to color-class IDs. Quotient filters work well with the Bentley-Saxe transformation because the quotient-filter merging algorithm consists of a linear scan over the inputs and the output and hence is I/O efficient.

## 3 Methods

In this section we describe the transformation of static Mantis to dynamic one using Bentley and Saxe (1980).

### 3.1 Dynamic Mantis

We make Mantis dynamic using the Bentley-Saxe (Bentley and Saxe, 1980) transformation, similar to LSM trees. Figure 1 shows the overall structure of our dynamic Mantis construction. The top level (level 0) is a collection of Squeakr (Pandey *et al*., 2017c) files. A Squeakr file is a representation of the *k*-mers present in a single sample. The other levels are all independent static Mantis indexes. Each level covers a disjoint set of samples. Thus, to determine the set of samples that contain some *k*-mer *x*, we query for *x* in each Squeakr file in level 0 and in each of the mantis indexes at the lower levels. The overall result is the union of the sets returned from each level.

**Fig. 1:**
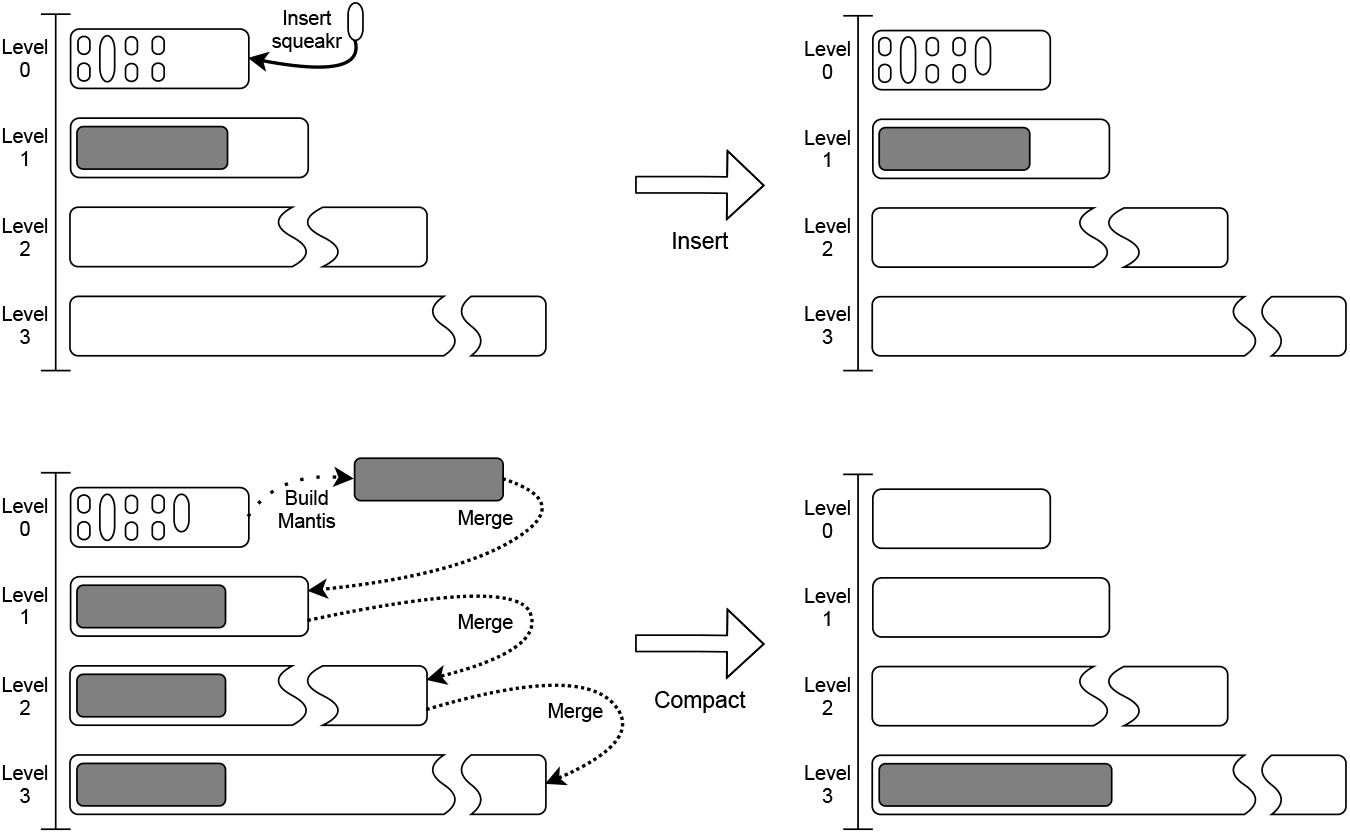
The two main operations in dynamic Mantis. For the *insert* operation, we only add a new Squeakr file to the list of Squeakrs in level 0. During the *Compact* operation, which is triggered whenever the number of Squeakrs passes the threshold for level 0, we first build a Mantis index over all the Squeakrs in level 0 and do a cascading merge of Mantis indexes until no level is full. Note: for graphical clarity, the levels are not drawn to scale.

To add a new raw sequencing sample to the index, we first use the *k*-mer counter Squeakr (Pandey *et al*., 2017c) to create a compact representation of all the *k*-mers in the sample (without counts). We then add this Squeakr file to level 0 of the Mantis index.

Since queries must examine every Squeakr file in level 0, we must not let level 0 grow without bound. Once level 0 reaches some threshold number of Squeakrs *M*, we perform a *compaction*. In the compaction step, a Mantis index is constructed from all Squeakr files in level 0 and the resulting Mantis index is merged into the first Mantis index at level 1. At this point, we can remove all the Squeakr files from level 0, since they are now covered by the Mantis index at level 1.

The Mantis indexes at each level are also limited in size—the index at level *i* is limited to size *r*^*i*^*M*, where *r* is a parameter that trades off between insertion and query performance, as described in Section 2. Whenever merging into level *i* causes the index on level *i* to exceed its maximum allowed size, then we merge level *i* into level *i*+1.

As in a cascade filter or LSM tree, the total number of levels is 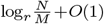, where N is the total number of samples in the index. The total cost of inserting all the samples is 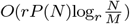, where P(N) is the cost of merging two indexes of size O(N) (see Bentley-Saxe Bentley and Saxe (1980) for details of the analysis).

Queries must query up toM Squeakrs in level 0, plus up to 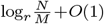 Mantises in the remaining levels. Querying for a k-mer in Squeakr takes O(1) time, so queries in our Dynamic Mantis cost 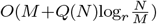, where Q(N) is the cost to query a Mantis index of N samples. So, for example, if we set r=2 andM=logN, then queries in our Dynamic Mantis are at most logarithmically more expensive than in a single Mantis index.

To support efficient compaction, we need to efficiently merge two Mantis indexes, i.e. we need to make P(N) as small as possible. In section 3.2, we describe what merging two Mantis indexes means at the schema level, i.e. without worrying about the low-level representation of the maps from k-mers to color-class IDs and from color-class IDs to colors. Later in section 3.3, we describe an efficient algorithm for merging two MST-based Mantis indexes. We further explore the particular challenge we face for having a memory-scalable merge in section 3.4.1 and provide a data-driven solution to address it.

### 3.2 Merging Mantis indexes

The Mantis merge algorithm takes as input two Mantis indexes for disjoint sets of samples *E*_1_ and *E*_2_ and produces an output Mantis index over *E*_1_⋃*E*_2_. Let *K*_1_ and *K*_2_ be the maps from k-mers to color-class IDs, and let *C*_1_ and *C*_2_ be the maps from color-class IDs to color classes (i.e bit-vectors). Each map *K*_*i*_ contains the unique set of *k*-mers across all the samples covered by Mantis index *i*. For simplicity, assume *K*_*i*_[*x*] = 0 for any *k*-mer *x* that is not in any sample in *E*_*i*_ and assume *C*_*i*_[0] is the all-zeros bit-vector with size equal to the total number of samples covered by the index.

Our task is to construct new maps *K* and *C* such that for every *k*-mer *x, C*[*K*[*x*]] = *C*_1_[*K*_1_[*x*]] ° *C*_2_[*K*_1_[*x*]], where ° represents bit-vector concatenation. We can view the two input indexes as mapping each *k*-mer *x* to a pair of color-class IDs, (*K*_1_[*x*],*K*_2_[*x*]) and each pair of color-class IDs (*c*_1_,*c*_2_) to a color class *C*_1_[*c*_1_]°*C*_2_[*c*_2_]. Thus we can break the process of constructing *K* and *C* into three steps. First, we construct a map *P* from color-class ID pairs to new color color-class IDs. We then construct *K*[*x*]=*P* (*K*_1_[*x*],*K*_2_[*x*]) for every *k*-mer *x*. Finally, we construct the new map *C*[*d*]=*C*_1_[*d*_1_]°*C*_2_[*d*_2_] for each mapping of the form *P* [*d*_1_,*d*_2_]=*d* in *P*.

### 3.3 Merging MST-based Mantis indexes

In this section, we explain the method for merging two MST-based Mantis indexes.

#### 3.3.1 Merging the *k*-mer maps

The first step is to construct the map *P* described above. This requires that we enumerate all the color-class ID pairs from the inputs *K*_1_ and *K*_2_.

The *k*-mer map in Mantis is implemented as a counting quotient filter (CQF) (Pandey *et al*., 2017a). A counting quotient filter stores the *k*-mers in order of their hash values, which means that we can efficiently enumerate all the mappings (*x,c*) in each input *K*_*i*_, in hash order. Since all Mantises use the same hash function, *K*_1_ and *K*_2_ are sorted in the same order, so we can use a variant of the merge algorithm from merge-sort to enumerate all the mappings (*x*,(*c*_1_,*c*_2_)), where *x* is a *k*-mer, *K*_1_[*x*]=*c*_1_ and *K*_2_[*x*]=*c*_2_. (Recall that if a *k*-mer *x* does not occur in one of the inputs, then we treat it as if it is assigned to color-class ID 0 in that map.)

##### Assigning new color-class IDs

A straightforward way to construct *P* would be to enumerate all the mappings (*x*,(*c*_1_,*c*_2_)) from the inputs *K*_1_ and *K*_2_, adding each color-class ID pair (*c*_1_,*c*_2_) to an in-memory hashtable. We could also use the hash table to record how often we see each color-class ID pair, so that we can then assign new color-class IDs in order of decreasing popularity (see Section 2 for how this saves space in the CQF).

However, the number of distinct color-class ID pairs can quickly grow to tens of billions, making this a RAM bottleneck. To skip the RAM overhead of this operation, we write all the pairs to disk in a file *F*, and then make the contents of the file unique. We also use sampling, as in the original Mantis paper Pandey *et al*. (2018), to construct a table *T* of the most popular color-class ID pairs. We assign the elements of *T* to new color-class IDs in descending order of popularity. For the remaining color-class ID pairs stored in *F*, we construct a minimal perfect hash function (MPHF) *H* mapping them to integers in the range 1,…,|*F* |. We then set *P* [*c*_1_,*c*_2_] to be *T* [*c*_1_,*c*_2_] if (*c*_1_,*c*_2_)∈ *T*, or |*T* |+*H*[*c*_1_,*c*_2_] if (*c*_1_,*c*_2_)∈ *T*. This ensures thatP assigns themost popular color-class IDpairs to lowcolor-class IDs.

##### Filling the output CQF

Given *P*, we can now construct *K*, the new quotient filter mapping *k*-mers to their new color-class IDs. We iterate over the input quotient filters *K*_1_ and *K*_2_ for the second time. This time, for each *k*-mer *x*, we insert (*x,P* [(*K*_1_[*x*],*K*_2_[*x*])]) into the output CQF.

##### Analysis

Let *n* be the sum of the *k*-mers in the two input indexes. Constructing *F* takes *O*(*n*) time, since we simply write at most one entry to *F* per *k*-mer in the input. Note that we don’t need to sort list of items in *F* completely—we need only uniquify it. Assuming we have, say, 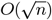 memory, we can partition *F* into 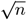 sub-files and route color-class ID pairs to partitions based on a hash function. We can then load each partition and insert all its elements into a hash table in memory and write out the results. Thus constructing *F* and uniquifying all its color-class ID pairs can be done in *O*(*n*) time. Constructing the MPHF on *F* also takes at most *O*(*n*) time (Limasset *et al*., 2017). Constructing *K* =*K*_1_∪*K*_2_ takes *O*(*n*) time since we can iterate over *K*_1_ and *K*_2_ in *O*(*n*) time and each lookup in *P* takes constant time. Furthermore, all the above steps either operate on in-memory data structures or can be accomplished by sequential scans of on-disk data, so the total I/O cost is *O*(*n/B*), where *B* is the block size of the storage device.

#### 3.3.2 Merging the color class representations

The high-level idea of the MST-merging algorithm is to simply run the MST algorithm used to construct an MST-based Mantis index from scratch. However, attempting this in a straightforward manner can result in an unacceptably slow merge algorithm; we take care to address these inefficiencies (see Section 3.3.3). Supplementary Figure 1 depicts the required modifications to the original MST construction process to convert it to merging two MSTs. We construct the new MST by constructing a graph on the set *C* of new color-class IDs. For each edge between *k*-mers *x*_1_ and *x*_2_ in the merged DBG, we add an edge between *K*[*x*_1_] and *K*[*x*_2_], where *K* is the merged map from *k*-mers to color-class IDs. The weight of the new edge is the Hamming distance between the color classes associated with color-class IDs *K*[*x*_1_] and *K*[*x*_2_], i.e. the edge weight is the Hamming distance between *C*_1_[*K*_1_[*x*_1_]]°*C*_2_[*K*_2_[*x*_1_]] and *C*_1_[*K*_1_[*x*_2_]]°*C*_2_[*K*_2_[*x*_2_]], where *C*_1_, *C*_2_, *K*_1_, and *K*_2_ are the input *k*-mer and color class maps. Once we have this graph, we can run the MST construction algorithm, completing the process of merging the Mantis indexes.

The main challenge in MST-based Mantis merge compared to MST-based Mantis construction is to efficiently compute the weight of each edge in this graph, which requires being able to efficiently query the input color-class tables *C*_1_ and *C*_2_. The original MST-construction algorithm takes as input the classic Mantis representation of *C*_1_ and *C*_2_, which supports constant-time queries. However, when merging MST-based color-class representations, we have access only to the MST representation of *C*_1_ and *C*_2_, and the MST-based representation does not support constant-time queries. Looking up these color-class IDs in the input MSTs requires walking the input MSTs from each queried ID to the root, applying the bit differences along each edge (Almodaresi *et al*., 2020). The lengths of these paths can be arbitrarily long (bounded only by the number of color classes), causing queries to be quite slow. We need to overcome this problem to ensure merging can be done efficiently.

#### 3.3.3 Static MST cache

We now describe an efficient pre-computation over the input MSTs that allows us to ensure that these color queries can be performed in constant time on average. During the precomputation step, we construct a static cache of nodes in the input MSTs. By precomputing and caching the bit-vectors for a subset of the color classes, queries can stop walking the MST whenever they reach one of the cached color classes. By choosing the cached nodes carefully, we can ensure that the average number of steps taken per query is constant.

One important advantage we have is that the color-class IDs to be queried are known ahead of time. We have a list of edges (as pairs of color-class IDs), so we know exactly what color-class ID pairs appear and how many times each color-class ID will be queried. We use this information to design a “static cache” that is populated once before the start of the query step and then used during the entire MST merge process. The cache can be further tuned for the lookup cost that the user wants to achieve, trading off memory for more efficient lookups.

The idea behind our computation is to maintain, for each node *x*, the number of queries *V* [*x*] that will visit that node, and the total distance *D*[*x*] walked by those queries. We start with *D* and *V* corresponding to having all nodes in the cache, and we discard nodes, from the leaves up, transferring the costs to their parents, until the average cost of all queries that visit a node exceeds some threshold *c*.

1. Initialize *D*[*x*] = 0 for every node in the MST.
2. Set *V* [*x*] to be the number of times *x* will be queried during MST construction. (This can be computed by walking over the DBG).
3. Set *L*={} (*L* will contain all the color-class IDs that should be loaded into the cache).
4. Start a post-order traversal on the MST. For each node *x*:
  a. Set *D*[*x*]=Σ_*d*∈children(*x*)_*V* [*d*](1+*D*[*d*]).
  b. Set *V* [*x*]=*V* [*x*]+Σ_*d*∈children(*x*)_*V* [*d*].
  c. If *c*−1 *<D*[*x*]*/V* [*x*], then add *x* to *L* and set *V* [*x*]=0.
5. Query the color-class IDs in *L* and put the color bit-vectors in the “static cache”. Since the list has been populated through a post-order walk, the ancestors of each node appear after the node in the list. Therefore, if we construct the color bit-vectors starting from last color-class ID to first in the list, we can even use the “static cache” while constructing it.

##### Analysis

The most expensive step of the algorithm is initializing *V* [*x*] by scanning the DBG. This takes *O*(*n*) time, where *n* is the number of *k*-mers in the DBG. The rest of the algorithm runs in time *O*(*e*), where *e* is the number of color classes, which is guaranteed to be less than or equal to *n*. The algorithm ensures that the average query walks a distance of at most *c* edges. Furthermore, the cache is guaranteed to contain at most a fraction 1*/c* of the nodes in the color class graph because (1) The average number of steps taken by queries that reach a node *x* in the cache is always larger than *c*−1 and less than or equal to *c*, so (2) there must be at least one query that traverses at least *c* edges (and hence *c*−1 non-cached nodes) before reaching *x*, so (3) there must be at least *c*−1 nodes that were not added to the cache for every node that was added to the cache. Thus *c* trades off cache space vs. query speed. Note that this is a worst-case analysis. As Supplementary Figure 2 shows, in practice, when merging Mantis indexes over RNA-sequencing samples we observe much smaller fractions than 1*/c* to achieve max distance of *c*.

### 3.4 Improving query and MST construction scalability

As reported in the original MST-based Mantis paper (Almodaresi *et al*., 2020), the CQF that maps *k*-mers to color-class IDs is the largest part of the Mantis index and, therefore, the CQF becomes a scaling bottleneck as we build larger indexes. The CQF becomes a bottleneck because both MST construction and Mantis queries require loading the entire CQF into RAM. In this section, we describe a method for dividing the CQF into smaller *partitions*, so that queries and MST construction need to load only a small number of partitions into RAM at any time. Note that the CQF partitions we consider here are not necessarily disjoint, and in fact commonly overlap.

We first describe the requirements that MST construction imposes on our partitioning scheme. During MST construction, Mantis constructs a graph on all the color-class IDs by iterating over all the edges of the DBG and adding a corresponding edge to the color-class graph. Mantis stores the DBG simply as a hash table (i.e. a quotient filter) of all the nodes, so iterating over the edges requires iterating over all the nodes and querying for every possible neighbor. Since the nodes are stored in a hash table, each neighbor query involves a random access to the table (Almodaresi *et al*., 2020). Once the table becomes too large to fit in RAM, the random accesses become a performance bottleneck.

We avoid loading the entire CQF into RAM by using minimizers (Roberts *et al*., 2004) to partition Mantis’s monolithic CQF into smaller CQFs. The key property we need is that all edges of the DBG connect vertices in the same partition. If we can achieve this property, then we can enumerate all the DBG edges by loading each partition one at a time and enumerating the edges within that partition.

We achieve this using the technique of Chikhi *et al*. (2016). The core idea is to partition the de Bruijn graph by minimizer. However, this is not a disjoint partitioning. Each *k*-mer is assigned two minimizers, one for its (*k*−1)-base prefix and one for its (*k*−1)-base suffix (for most *k*-mers, these will be the same minimizer). Thus all adjacent vertices will have at least one minimizer in common. By placing each *k*-mer into possibly two different partitions (one for each of its minimizers), we can ensure that all edges in the DBG connect vertices that are present in the same partitions. Note that this idea can be used with any of the common definitions of minimizer (e.g. lexically first, smallest hash value, etc.).

One could, in theory, create a separate CQF corresponding to each minimizer, but this approach has two main issues: (1) it requires storing 4 CQF files on disk for partitioning based on a minimizer of length ; and (2) it may lead to CQFs with very small numbers of *k*-mers, which leads to small CQFs with few slots and large remainders. This would therefore result in using the CQF in its most inefficient way. To overcome these issues, we greedily place the *k*-mers for multiple consecutive minimizers into the same CQF until the CQF reaches a minimum size, *U*. We then store the minimizer boundaries between partitions so that we can direct queries for *k*-mers to the appropriate CQF.

#### 3.4.1 Merging partitioned CQFs

Partitioned CQFs make merging the CQFs slightly more complicated, primarily because the two inputs might be partitioned differently. If the two input partitioned CQFs had the exact same boundaries, then each CQF would cover the exact same set of minimizers. We could therefore just iterate over the CQFs in the partitioned CQF, merging each pair of corresponding CQFs using the standard CQF merging algorithm.

When the two input partitioned CQFs have different boundaries, then we are not guaranteed to see all the *k*-mers in the same order if we iterate over the two inputs. For example, if the first CQF of the first input includes all the *k*-mers with minimizer “AAAAA”, but the first CQF of the second input does not include *k*-mers with minimizer “AAAAA”, then we cannot simply merge the first CQFs in each input, since some *k*-mers will be missing from the second input’s first CQF.

We solve this problem by buffering *k*-mers by minimizer until we are guaranteed to have seen all instances of a given *k*-mer from all input CQFs. More concretely, let *K*_1_,…,*K*_*a*_ and 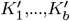 be the CQFs for each input, and let *τ*(*K*) be the smallest minimizer covered by CQF *K*. We first construct a list *L* of all the CQFs (from both inputs) sorted by *τ*. We then construct an array *S*, indexed by minimizer, of *k*-mer sets, where each set is initially empty. We then iterate over all the CQFs in *L*, adding all their *k*-mers to each *k*-mer’s minimizer’s entry in *S*. When a *k*-mer has more than one minimizer, we consider only its minimizers that fall within the range of the current CQF. After processing *L*[*i*], we are guaranteed to have seen all *k*-mers with minimizer up to *τ*(*L*[*i*+1]). Thus all the *k*-mers in *S* with minimizer less than *τ*(*L*[*i*+1]) are eligible for output. Once we have accumulated *U k*-mers in the sets of *S* indexed by minimizers less than *τ*(*L*[*i*+1]), we dump them all into an output CQF and reset those entries in *S* to empty sets, freeing up that memory.

The entire process is depicted in Supplementary Figure 3. In addition, we provide the pseudo-code for the merge operation in Supplementary Algorithm 1. We also discuss the details of the memory management and parallelization of the merging procedure in Supplementary Section 3.

##### Analysis

This algorithm for merging the CQFs in a Mantis index runs in linear time, just like the standard algorithm for merging two CQFs. This is because we can use hash tables (or even just linked lists) to implement the *k*-mer sets in *S*, so that adding each *k*-mer to its entry in *S* takes constant time. To build each output CQF, we just iterate over the *k*-mer sets in *S* that are to be included in the output, inserting their *k*-mers into the output CQF. Since the output CQF is a hash table, insertions take *O*(1) expected time. Thus we perform *O*(1) total work for each *k*-mer, for a total merging cost of *O*(*n*).

## 4 Results

Our evaluation is intended to answer the following questions:

- How well does mergeable Mantis scale in comparison to other state-of-the-art exact colored-DBG representations, such as VariMerge and Bifrost, in terms of time, memory, and intermediate disk space?
- How does the partitioned CQF affect memory and speed of queries in Mantis?
- How expensive, in terms of time, memory, and storage, are updates and queries in our dynamic Mantis construction? These are the essential questions for understanding how well Mantis will scale to huge data sets in terms of construction, queries, and updates.

### 4.1 System Specification and Input Data

#### 4.1.1 System Specifications

As with prior work (Almodaresi *et al*., 2019, 2020; Pandey *et al*., 2018), the input to Mantis is a list of *Squeakr* files (Pandey *et al*., 2017c). The Squeakrs are each constructed from a specific FASTQ file selected from human RNA short read sequences publicly available via SRA (Leinonen *et al*., 2010). The system for all the experiments is an Intel(R) Xeon(R) CPU (E5-2699 v4 @2.20GHz with 44 cores and 56MB L3 cache) with 512GB RAM and 4TB TOSHIBA MG03ACA4 ATA HDDs running Ubuntu 16.10 (Linux kernel 4.8.0-59-generic). All Mantis experiments use the MST-based color-class table representation. Partitioned CQFs target 2.4GB per CQF. Partitioned CQFs target 2.5GB per CQF limit. As per our procedure for filling these CQFs, we can either insert all of the *k*-mers with the same minimizer value into a partitioned CQF or none if the set of *k*-mers passes the partitioned CQF threshold (a minimizer cannot be split over multiple partitions). We choose 2.5GB as the smallest size for a partitioned CQF, since it is sufficiently large compared to the average number of *k*-mers sharing a minimizer (using a random minimizer order) to allow most CQFs to be near full.

Constructing and merging Mantis are both conducted using 16 threads, as is constructing Bifrost. VariMerge (Muggli *et al*., 2019), however, supports only single-threaded construction. Query benchmarks are performed using a single thread for Mantis and 16 threads for Bifrost. VARI requires users to specifyn the maximum number of samples at *compile time*, and this parameter affects the memory usage and size of the resulting indexes. In order to show VARI in the best possible light, we always use VARI executables that were compiled with the smallest possible limit. Time and memory are evaluated using “/usr/bin/time” command for all the experiments. Elapsed time is reported for time and maxRSS is reported for peak memory (which we call memory throughout the paper). Commands used to run each of the experiments and the options used for different tools are listed in Section 4

#### 4.1.2 Input Data

For the input to our experiments, we download 39,400 FASTQ files from NCBI (O’Leary *et al*., 2016). The list of accessions is available in the project github repository^1^. For our core experiments we constructed the Squeakr files in advance for all the samples over roughly four weeks on a cluster of 150 machines with a 24-disk array connected via a 10Gbps link. For each file, only the *k*-mers with abundance value more than a predefined threshold are selected. The value of the threshold is decided based on the size of the gzipped FASTQ file. This prefiltering step is useful to eliminate spurious *k*-mers that occur with insignificant abundance and has been adopted from the original SBT paper Solomon and Kingsford (2016b). The *k* chosen for the *k*-mers across all the experiments is fixed at *k* =23. The total space required to store all FASTQ files and their corresponding Squeakr files is 2.33TB and 970GB, respectively. Our Bifrost comparison experiments used a different approach explained in Section 4.2.

### 4.2 Comparing Mantis with Bifrost

Figure 2 compares Mantis and Bifrost in terms of construction time and memory, and the size of the resulting index. Note that Bifrost does not support merging, so we just compare its construction performance with Mantis’ construction performance.^2^ We run Bifrost in its color-based indexing mode. Bifrost differs from Mantis in two important ways that we must take into account in order to make a fair comparison. First, Bifrost builds its index directly from FASTQ files^3^, whereas Mantis builds its index from Squeakr files (which are built from fastq files). Thus, for fairness, we include the Squeakr construction time in Mantis’s build time. Second, Bifrost filters all *k*-mers that occur only once across all input fastq files. Mantis performs filtering independently on each input file as part of the Squeakr building process. Thus Mantis may filter out more *k*-mers than Bifrost (i.e. a *k*-mer that occurs once in each input would be filtered by Mantis but not Bifrost). To provide a meaningful comparison, we compare Mantis performance in two conditions. In the first condition, we include all *k*-mers in the samples, i.e. we perform no filtering at all. In the second condition, we perform Mantis’ standard per-file singleton filtering.

**Fig. 2:**
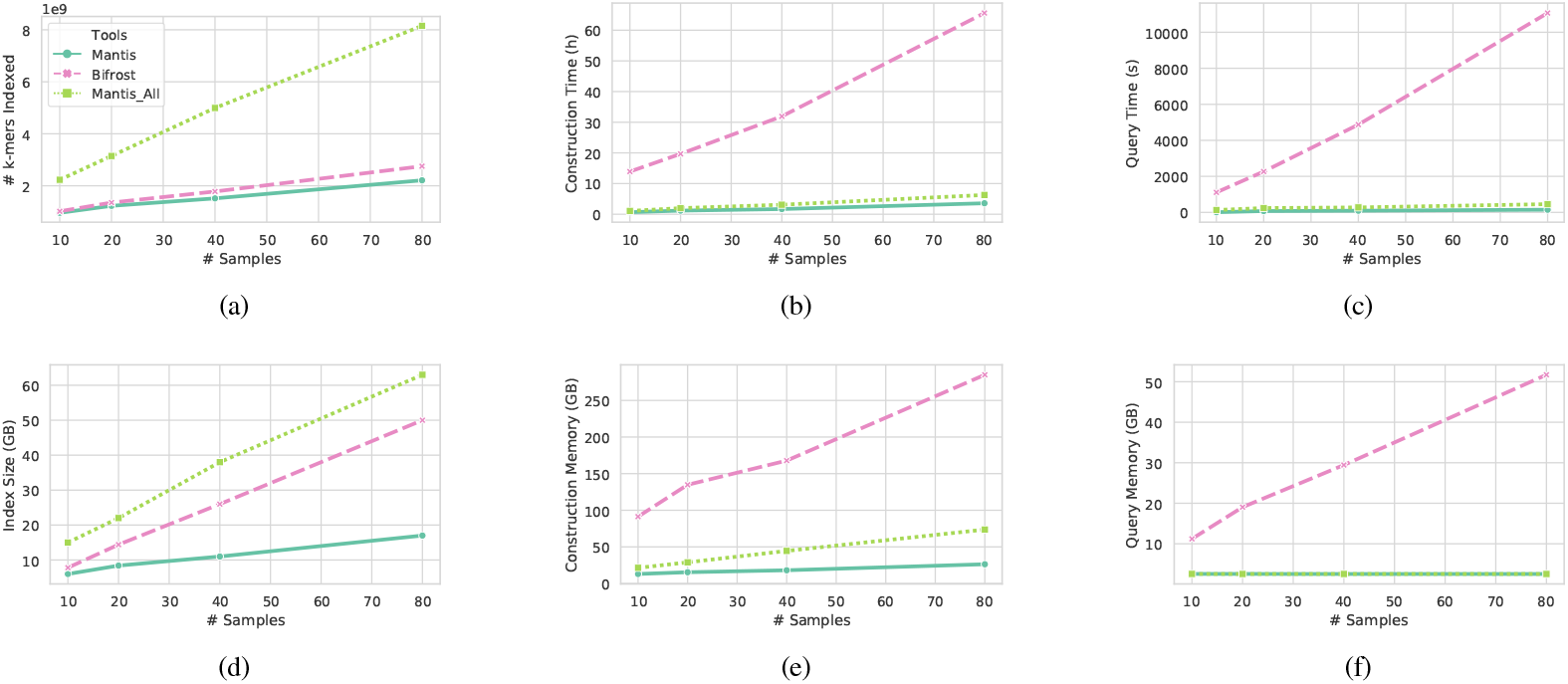
Comparing the construction performance, query performance, and index size of Bifrost and Mantis. In constructing Mantis, singleton k-mers per file are filtered out; Mantis_All includes all the k-mers; and Bifrost filters any k-mer that occurs once across all the input files in each experiment (10,20,40,80). Mantis peak memory is evaluated over the full pipeline including constructing Squeakrs. Query benchmarks are over 1000 randomly chosen human transcripts as input query sequences

We construct Mantis and Bifrost indexes over 10, 20, 40, and 80 samples to compare the scalability of the tools using 16 threads for both. Mantis always runs about an order-of-magnitude faster than Bifrost (Figure 2b) and uses about an order-of-magnitude less memory (Figure 2e), even when we turn off singleton filtering.

Figure 2a shows that, despite the slight difference in filtering rules, the total number of *k*-mers indexed by Bifrost and Mantis are quite close, so we can make reasonable comparisons between the sizes of the resulting indexes. The resulting Mantis indexes (filtering singletons) are about half the size of Bifrost’s indexes. Even when we include all *k*-mers in the Mantis index (4× more *k*-mers than Bifrost’s), Mantis’ index size is only ∼1.2× larger than Bifrost’s.

Mantis queries are orders-of-magnitude faster than Bifrost (Figure 2c) while also requiring an order-of-magnitude less memory (Figure 2f). Specifically, Mantis queries in the filtered case are 74× faster than Bifrost and Mantis queries on the unfiltered index are 24× faster than Bifrost’s queries on its filtered index. Mantis’s memory is a constant ≈ 2.5GB (which is equal to the size of one CQF partition). Mantis queries are single threaded but we ran Bifrost queries using 16 threads. We believe the gap between the query performances is largely due to the fact that Bifrost records the compacted dBG as a GFA file, and does not record its index on disk. Hence, for each query call, Bifrost must first reconstruct an index over the graph from the GFA file and the color encoding.

Because we are comparing Mantis’s two-step process (Squeakr plus Mantis) to Bifrost’s single-step construction mode, Mantis uses some intermediate disk space for Squeakr files, whereas Bifrost does not use any intermediate space. However, in our experiments, the peak space used by both systems was basically the same. For example, in our 80-sample experiments, the total space for all the Squeakr files plus the Mantis index was 20% smaller than Bifrost’s index (42GB vs. 50GB). However, Mantis indexed about 80% as many *k*-mers as Bifrost in this experiment (see filtering discussion above), so the peak space per indexed *k*-mer is basically the same in both systems.

Bifrost’s construction performance in our benchmarks is very different from the performance reported in the Bifrost paper Holley and Melsted (2020). We believe this difference is due to differences in the heterogeneity of the input samples. Bifrost is a compacted de Bruijn graph, which means that Bifrost attempts to save space by storing non-branching paths as a single sequence, rather than a collection of overlapping *k*-mers. This works well when the samples being indexed are very similar, so that the colored DBG contains many long non-branching paths. However, when the input samples are more diverse, the average non-branching-path length goes down until, in the extreme, each non-branching path is only a handful of *k*-mers long. In this case, there is little space to be saved from the compacted representation, but the index still pays for the complexity of the encoding. This is why Bifrost shows great scalability when indexing a set of very similar sequences such as Salmonella genomes but much worse performance when dealing with greater variation and diversity in the database, as in these sequencing samples.

### 4.3 Comparing Mantis with VariMerge

The target in this section is to evaluate the merging performance of Mantis and compare it with other tools that support the same operation. Therefore, we do not compare the input index preparation steps (i.e. *k*-mer counting and index construction)^4^. We evaluate the memory and time required to merge Mantis indexes, and the sizes of the resulting indexes and compare with VariMerge which is, to our knowledge, the only other exact de Bruijn graph representation that supports a “merge” operation.

Note that the merges in our experiments were not always balanced. Specifically, to construct the 500-sample indexes, we merged indexes on 200 and 300 samples. Similarly, the 5000-sample index was constructed by merging indexes of 2000 and 3000 samples. The rest of the indexes were constructed from balanced inputs, e.g. the 10,000-sample index was constructed by merging two 5,000-sample indexes.

The scaling plots in Figure 3 show that the Mantis merge operation uses less than a quarter the memory and a third the time of VariMerge. Furthermore, Mantis indexes are less than half the size of VariMerge’s indexes. In fact, VariMerge ran out of memory while attempting to merge two indexes of 5K samples into a 10K index on our system, which has 512GB of RAM.

**Fig. 3:**
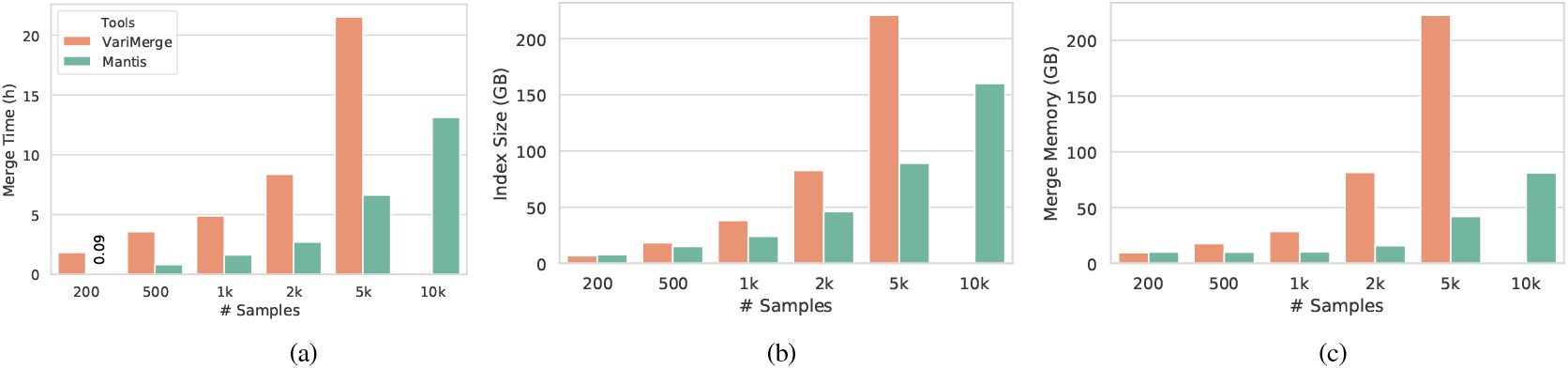
Benchmarking VariMerge and Mantis merge for building different number of samples from 200 up to 10k. The values on the x axis show the size of the output. The results show Mantis merge is superior in all the metrics.

### 4.4 Partitioned CQF Query Benchmarks

We measure the impact of partitioned CQFs on the query time and memory requirement of Mantis by performing bulk queries on two Mantis indexes of 10K samples, one of which uses partitioned CQFs and the other uses a monolithic CQF. As the plots in Figure 4 show, the query time with partitioned CQFs is very similar to that of MST-based Mantis with a single CQF, while the memory required is up to 7× less. Although, the partitioned CQF introduces some overheads when performing small batch queries, these overheads become insignificant in large batches. We also note that, if we load all the partitions into memory, then the partitioned CQF version is just as fast as the original version of Mantis, but this eliminates the memory savings of partitioned CQFs.

**Fig. 4:**
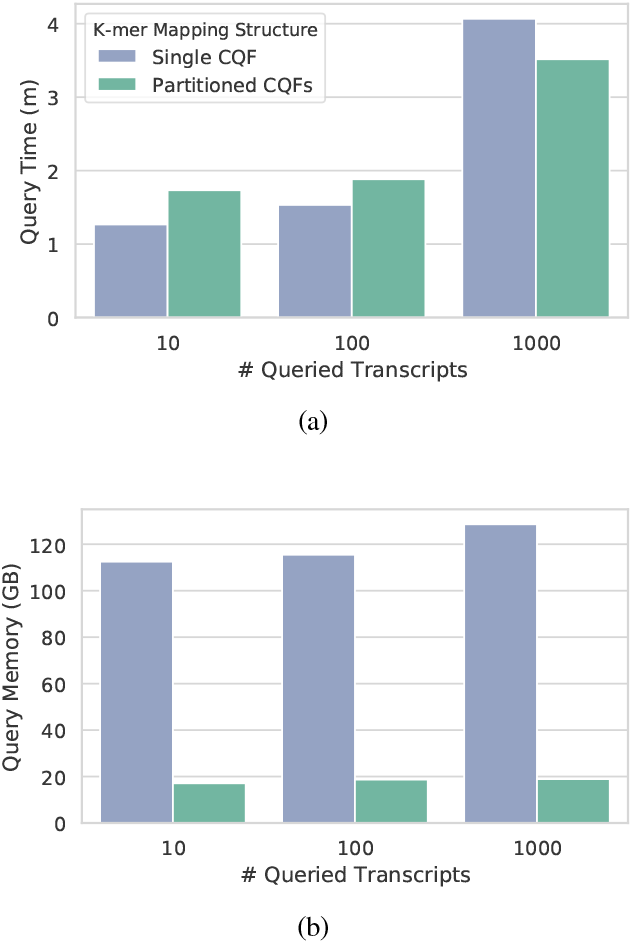
Time and memory requirements of bulk queries in Mantis with partitioned CQFs versus Mantis with a single CQF.

### 4.5 Dynamic Mantis Benchmarks

The purpose of this experiment is to explore the scalability and updatability limits of our Dynamic Mantis index. No other indexing scheme that we evaluated could index more than 10k experiments, so we cannot make comparisons to other schemes at this scale. Rather, our goal is to understand the performance and limitations of Dynamic Mantis.

In our Dynamic Mantis benchmarks, level 0 accumulated up to *M* = 100 Squeakr files before being merged into level 1, and we used a growth factor of *r* = 4 between levels. We measure the size of the Mantis levels (i.e. level 1 and up) based on the number of CQFs in the partitioned CQF. Level 0 was considered full when it exceeded 5 CQF partitions. So level 2 could have up to 20 partitions, level 3 could grow to 80, and so on. We start with an index of 10k samples, and add more samples in batches of 100.

Figure 5 shows the memory consumption and time for batch insertion as a function of number of samples. As the results show we observe a steady behavior in insertion time for the first 4000 samples after 10k. We observe a close to constant insertion time for most of the batches of 100 samples and there is a regular peak of longer time (∼20,000 seconds) about every 500 samples. That is when we need to continue the merge more than 1 level up to level 2. There are three big peaks for those samples that trigger cascading merge up to level 3.

The trend in the insertion memory plot is similar to the time plot with close to constant memory for most of the insertions except for the ones that require big merges during the cascading merge process. At those points the memory increases linearly to the total size of the largest of the indexes being merged. The size of the index is not strictly increasing because merging can reduce redundancy from repeated *k*-mers across levels. One sample of such behavior is shown in figure 5d, where the size of the index for 32*k* samples is larger than size of the index after adding 7K more samples.

**Fig. 5:**
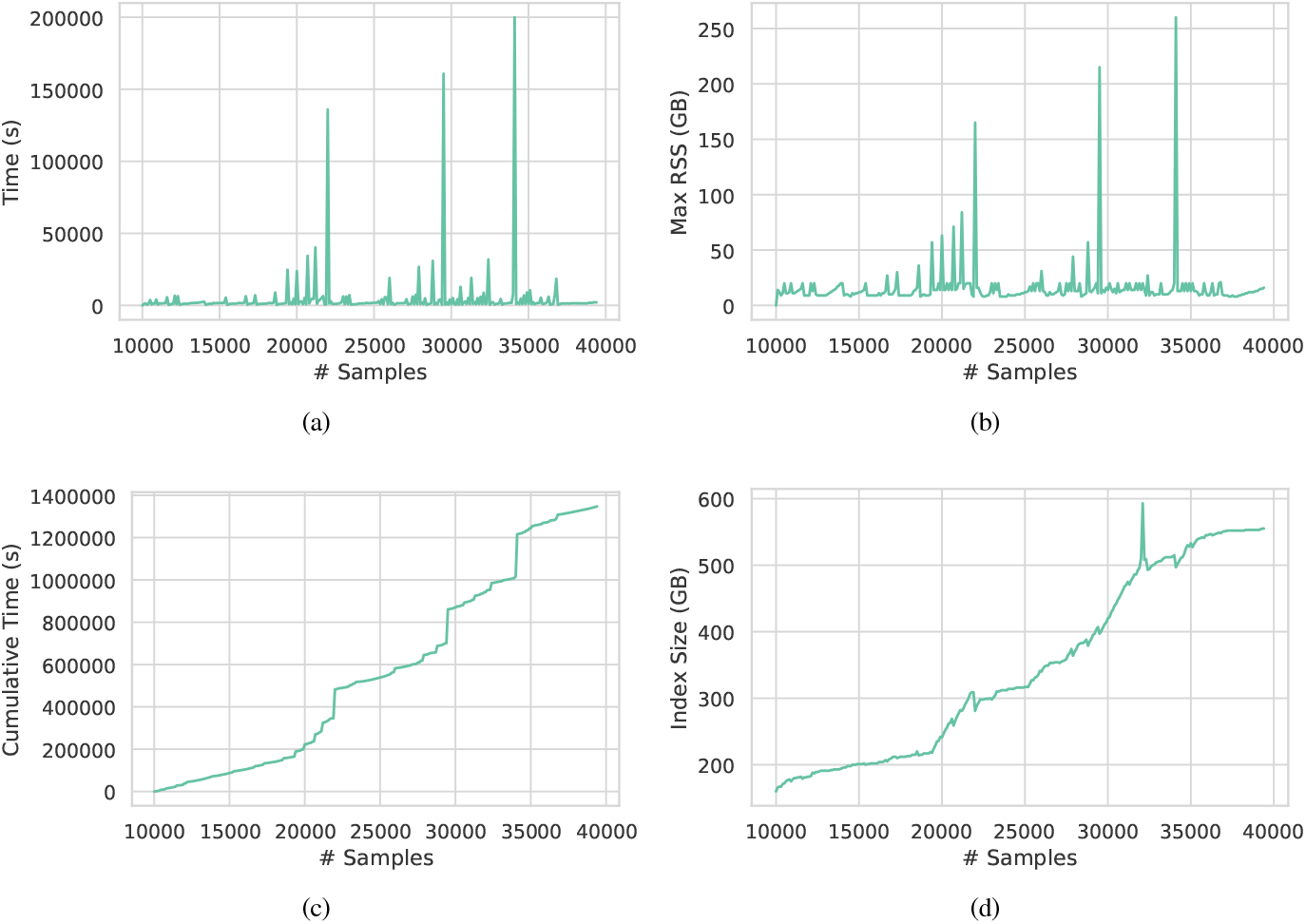
Performance of the Dynamic Mantis update process. The spikes in time and memory happen when the cascading merge happens with deeper and thus larger indexes.

Based on these results, we believe the main bottleneck to scaling Mantis to larger data sets will be the spikes in memory usage during merges. These memory spikes are primarily for loading the input MSTs. For example, during the memory spike at about 29,000 samples in Figure 5, the two MSTs being merged consumed over 100GB of RAM (even though they are stored in a compressed form in RAM), and the caches consumed an additional 40GB of RAM. Thus the MSTs and their caches accounted for over half the peak RAM. Our current merging algorithms require essentially random access to the input MSTs, so they need to be able to store the entire MSTs in RAM to get good performance.As a result, our algorithms use memory that is linear in the size of the MSTs. Future work on scaling Mantis should likely focus on sublinear-memory algorithms for merging MSTs.

We also benchmark the query time over the final index. Querying 1000 random human transcripts in an index of 39.5K samples took 1 hour and 48 minutes, including the index loading time.

## 5 Conclusion

In this work, we describe an incrementally-updatable sequence search index that maintains many of the favorable properties of the previous index, MST-based Mantis, such as being exact and providing fast bulk queries, but also advances the index with respect to its scalability and memory requirements. By incorporating the Mantis index into an LSM tree structure, we enable insertion of new samples without requiring reconstruction of the index. We provide a memory-efficient and highly-parallelized algorithm for merging two (partitioned CQF-based) MST-based Mantis inputs.

We also replace the CQF filter in Mantis with a collection of partitioned CQFs that are (individually) much smaller in size and can be loaded individually during the MST construction and query processes. This reduces the query memory requirements significantly and ensures that the CQF data structure is no longer the memory bottleneck in MST construction and merging. We also provide benchmarks for incremental updating of our data structure using Dynamic Mantis to show the amortized cost of adding new samples to our data structure. Using this procedure, we were able to construct a Dynamic Mantis index over more than 39K raw sequencing samples which takes only 410GB of space and can be easily extended to larger number of samples.

We believe that, with the current strategy, we are approaching the limits of the number of samples that are practical to index exactly on individual mid-range servers. However, it is relatively natural to consider scaling the Mantis data structure out. In such a scheme, disjoint collections of samples would be distributed among a network of machines, with each node maintaining a distinct Mantis index. Queries can then be broadcast and executed in parallel at each node, and collated upon return. As future work, we plan to explore the efficient construction, maintenance and query of such a distributed index that would allow indexing data at the scale of all unrestricted human RNA-seq samples and beyond.

## Footnotes

1 https://github.com/splatlab/mantis/blob/mergeMSTs/experiments/humanRNA_40k.accessions

2 Bifrost does support an update command for adding new FASTQ files to an existing index, but we found its performance in this data was substantially worse indexes are less than half the size of VariMerge’s indexes. In fact, VariMerge ran out of memory while attempting to merge two indexes of 5K samples into a 10K index on our system, which has 512GB of RAM. than constructing the index all at once, so we report only bulk construction results in this paper.

3 Bifrost does support input directly from GFA in addition to FASTQ files. This means that one can develop an approach in which FASTQ files are first processed into GFA files and then Bifrost builds an index on the GFA files directly. However, we found that, in practice, this approach took even longer than directly constructing Bifrost over FASTQ files. Thus we report only the single-step construction results (building the Bifrost index directly from the FASTQ inputs, in reference mode) in this paper.

4 Prior work has shown than Squeakr, the preprocessing tool for Mantis, is substantially faster than KMC2, the preprocessing tool for Vari and VariMerge (Pandey *et al*., 2017c). Our experience was consistent with those results. We also found that Mantis was generally faster than Vari in several preliminary index construction tests. For example, Mantis never took more than 5 hours to construct an index on 1000 samples, whereas Vari took anywhere from 8 to 23 hours. However, Mantis used 16 threads whereas Vari is single-threaded, so these results are not intended to be a systematic, apples-to-apples comparison.

