## Supplementary Material for "An Incrementally Updatable and Scalable System for Large-Scale Sequence Search using LSM Trees"

### 1 Merging Classic Mantis

The color class matrix is partitioned and stored as fixed-size blocks of color classes to the disk, instead of storing the table as a whole. This facilitates in keeping the working memory low, as a color class is not required to be present during the full lifetime of the Mantis merge algorithm. Let the input indices $cdbg_1$ and $cdbg_2$ have $d_1$ and $d_2$ partitions in the color class table. For simplicity, we assume color class partitions to be $1 - based$ indexed. We initialize $(d_1 + 1) \times (d_2 + 1)$ files on disk (called disk-buckets) to temporarily store color-ID pairs of $k$-mers. A disk-bucket $b_{i,j}$ with $i, j > 0$ contains the color-ID pairs $(id_1, id_2)$ where $id_1$ belongs to color class partition $i$ and $id_2$ belongs to color class partition $j$. In the case the $k$-mer is not present in one of the inputs, the color-ID pair would end up in a disk-bucket with $i = 0$ or $j = 0$. Disk-bucket $b_{0,0}$ is empty as it is, by definition, associated to the color-ID pairs where the associated $k$-mer is neither present in $cdbg_1$, nor in $cdbg_2$.

Since the CQFs of the indices $cdbg_1$ and $cdbg_2$ contain the hash-values of the $k$-mers in sorted order, we make simultaneous linear scans over the two CQFs for the distinct $k$-mers of the union CQF. For each distinct $k$-mer *key* of the union CQF, let its color-IDs be $id_1$ and $id_2$ from $cdbg_1$ and $cdbg_2$. If $id_1$'s color class is in partition $x$ and $id_2$'s color class is in partition $y$, then we store the pair $(id_1, id_2)$ to bucket $b_{x,y}$, with possible repetitions of the pairs. Having completed the union process, we filter out the unique $k$-mers at each bucket (by sorting and discarding duplicates of the $k$-mer hashes in the buckets).

Simply following this scheme to collect the distinct pairs results in too much repetition of pairs and, thus, a high memory demand. Since we store a color-pair per $k$-mer, the disk-buckets will cumulatively contain exactly $n_k$ pairs if the merged CDBG has $n_k$ $k$-mers, whereas the number of color classes is actually much smaller. In reality, not only can multiple $k$-mers can share the same color, but also the majority of $k$-mers actually belong to a small subset of colors, less than 1% of total colors [Pandey *et al.*(2018)Pandey, Almodaresi, Bender, Ferdman, Johnson, and Patro] in a typical dataset. The original Mantis data structure leverages this highly skewed abundance distribution of color classes by sorting the color classes based on their abundance values, and assigning IDs in reverse order of the abundance — *at least* for the colors observed in a sample subset. Since in the final space required to store each color-ID correlates with the ID value itself, assigning smaller

IDs to more abundant colors results in smaller size representation of the color-IDs. This utilization of the highly-skewed color distribution is popular in many other tools in the field as well [Almodaresi *et al.*(2017)Almodaresi, Pandey, and Patro, Holley and Melsted(2019)Holley and Melsted, Almodaresi *et al.*(2018)Almodaresi, Sarkar, Srivastava, and Patro]. We perform a sampling phase in which we analyze the color class abundance distribution of a subset of the $k$-mers. We sort the color class based on their abundances in the sample set, and keep a fixed number of the (approximately) most abundant pairs in a hash-map $H$ which maps a color-ID pair to its abundance. Given the uniform-randomness property of the hash function used to hash $k$-mers, in expectation, we will see the most abundant color classes in the first few million $k$-mers. After this sampling phase, we make the simultaneous linear scan over the CQFs and for each color-ID pair $(id_1, id_2)$ of a $k$-mer, we only add the pair to its corresponding bucket if it is not present at $H$.

### 2 Improving Memory and Performance of MST Merge

**Memory management:** Although encoding the color class representation in Classic Mantis using a MST in MST-based Mantis improves the index scalability, both in terms of query memory and disk usage, still the high memory consumption during MST construction remains a bottleneck for scaling the index to more samples. Almost all the steps of constructing the MST are memory-expensive if we prohibit intermediate disk usage. During the construction process, first, we construct a sparse graph of colors by only adding a subset of edges we believe are potentially low-weight edges. There we only add an edge between two colors if their corresponding $k$-mers are neighbors in the CDBG. This idea is based on the observation that neighboring $k$-mers in a CDBG tend to have similar colors. Practically, the stated heuristic helps reduce the order of the color-graph edges from $\mathcal{O}(n^2)$ down to $\mathcal{O}(n)$. This change in order, makes the whole MST construction practical in the first place. However, due to the large number of colors, $n$, the pipeline is still memory-hungry if implemented naively. We explain below a number of optimizations we adopt to improve the construction of the MST:

- *Store serially-accessed structures on disk.* Large structures, such as the color-graph edges, which are accessed once through serialized scanning can be stored on disk. The tradeoff of random access memory for time (and external memory) in such cases seems a practical way to improve scalability.

- *Discard edges with weight greater than a global threshold during edge weight calculation for the color-graph.* The input list of edges to Kruskal's algorithm [Horowitz and Sahni(1978)Horowitz and Sahni], which is used to construct the MST, should be sorted based on weights. Since the weight value is bounded above by the number of samples in the index (which we know ahead of time), during the weight calculation process, we perform a bucket sort on-the-fly, moving the edges into the corresponding weight bucket, which is a file on disk. We only store edges with a weight up to a predefined threshold based on the heuristic that, eventually, most of the high-weight edges would be discarded during MST construction. This decision will not affect the functionality of the MST construction, since the presence of the dummy node ensures a connected spanning tree can always be built. Thresholding the input edge weights can only lead to a sub-optimal spanning tree compared to one constructed on the full set of color-graph edges. Practically, we observed only a small effect of this heuristic on the size of the final MST.

- *Store dummy-edges in a different data structure than non-dummies.* As per our definition, one end of a dummy edge is always the zero vector, and as a result, for each dummy edge of $\langle c_i, c_{dummy} \rangle$, the weight is the number of set bits in $c_i$, and the edge delta is the indices of the set bits in $c_i$. Hence, we can store these edges, which are a considerable fraction of edges in a data structure specialized for dummies, and keep them always in memory. The structure to store weights is simply a bit-packed int-vector with $n$ slots of width $\lceil \log_2 n \rceil$.

- *Design a memory-efficient structure to store the weighted adjacency list for the MST.* The result of Kruskal's algorithm is the list of edges in the MST, and each edge's weight, in the form of an adjacency list. The required space to store the adjacency list is linear in the number of nodes (or edges) in the MST (i.e. number of colors in the CDBG). This can be quite costly if implemented naiëvely. We adopt an efficient representation of the weighted adjacency list that is explained in 2.1.

- *Fill parent vector via a hybrid DFS-BFS walk.* To fill out the parent vector from the adjacency list, we need to perform a DFS or BFS over the tree starting from the root. However, because the MST is large both in width and height when constructed over many samples, both BFS and DFS traversals require substantial memory to perform. A BFS or iterative DFS procedure require a lot of memory for book-keeping, and a recursive DFS would fail as a result of stack overflow. Instead, we use a hybrid traversal to fill out the parent vector. The idea is similar to iterative deepening search [Reinefeld and Marsland(1994)Reinefeld and Marsland], keeping the memory constraint in mind. We set a depth limit for DFS. We start a DFS from the root and set the parent-child relationship in parent vector. Every time we reach our the limit, we keep the list of nodes at that limit (similar to the bookkeeping in BFS, but only in one level), and restart a DFS from each node at the level up to the next level where the difference again passes the limit. We continue this, iteratively, until we observe all the nodes in the tree and complete filling the parent vector. Rather than storing the actual IDs of the nodes each time we stop the DFS, we keep a bit-vector of size $n$ (the total number of colors/nodes in the tree) and set the related bits for the nodes that are going to be the root for the next DFS start. In every iteration of the iterative DFS, starting from node $i$, we reset bit $i$ in the bit-vector. We find the set bits in each round by traversing the bit-vector. This DFS procedure can also be parallelized.

**Parallelization:** Given the scale of the data structures being constructed, it is practically important that construction be parallelized even if the construction procedure itself is designed to be efficient. In our implementation, we have kept this in mind, and the following steps of the algorithm have been parallelized:

- *Walking the partitioned CQFs to construct the edges.* If we assume that we have $t$ threads, each partitioned CQF is divided into $t$ equal-size parts. Each thread walks over the $k$-mers in its designated part of the partitioned CQF searching for the neighbors of the $k$-mer in the entire partitioned CQF. The edges are stored on disk to keep working memory low. Therefore, we make use of a multi-producer queue to store batches of edges on disk every often.

- *Calculating edge weights.* We simply divide the edges between threads.

Each thread is responsible for calculating the weights of the assigned subset of edges. For this, each thread needs to access the MST structure, and the associated structures mainly the static cache. The static cache is constructed only once before the start of this step and will not be modified at any subsequent point, so it can be safely shared across threads to read from. When storing these edge weights (temporarily) to disk, we again need to buffer the edges and then flush them to disk in batches. Practically, we have observed considerable unevenness in the distribution of edge weights. The vast majority of edges have small weight values and the weight distribution is highly-skewed. For example, for an index over $2k$ samples with close to 1 billion edges, less than $200,000$ of them had a weight ¿ 1000. To keep the balance in frequency of flushing the buffers for each weight bucket, we used a geometric distribution (common ratio $= \frac{1}{2}$) of the buffer sizes where the files assigned to smaller weights have larger buffers that take up more of the allocated space in RAM.

- *Filling Delta and Boundary vectors.* Although the process of querying the input MSTs is the same as calculating the weights, there is an additional complexity filling the delta vector. The size of the delta vector and the order of the deltas are already predetermined; the size is total weight of the MST and the order is the same as the edges in parent vector. This means that each thread can be assigned a start and end of the range it is allowed to fill in the delta vector. The goal is achieved by performing one extra pass over the parent vector to sum up the weights of all the edges belonging to the same thread segment, assuming each thread is responsible for extracting deltas of edges in a consequent section of the parent vector.

#### 2.1 Detailed design of the memory-efficient structure to store the weighted adjacency list for the MST

The MST merge process is highly memory-intensive. We pointed out some of the optimizations we consider during the implementation to improve memory consumption, one of which was using a succinct representation of the weighted adjacency list. Here we explain this idea in more details.

By the end of the MST finding process, we would have a list of selected edges and their weights. As shown in figure 1, the next step is assigning

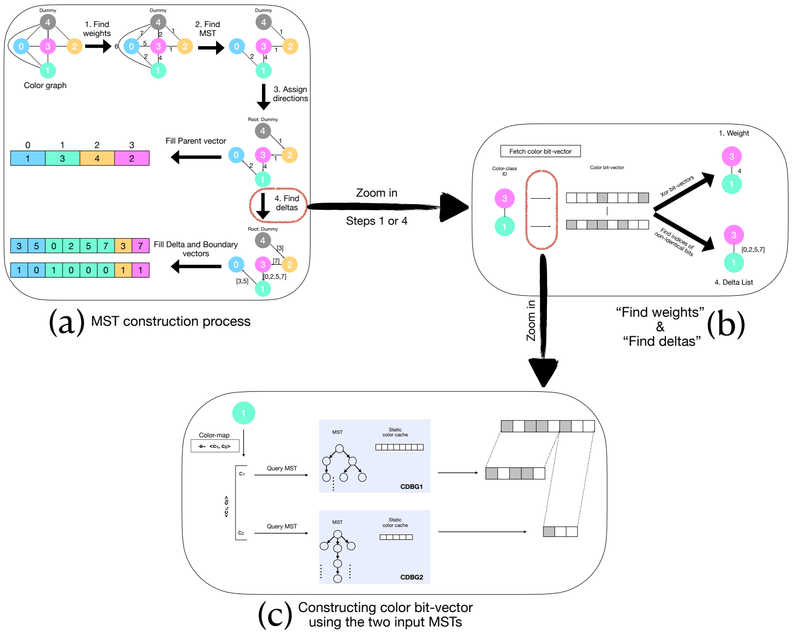


Figure 1: The full MST construction process (a) which is zoomed in two levels down in (b) and (c). The process in (a) basically follows the same steps as for MST construction in the MST-based Mantis index. The main point of difference is reconstruction of the color bit-vectors by querying the two input MSTs as shown in (c) rather than fetching the vector from the color class matrix. During the MST construction, step (c) will be repeated for each end of the edge in (b) and eventually for all the edges in the original color-graph (step 1) and later the final MST (step 4) in (a). Consequently, the main challenge is to make the color bit-vector reconstruction process as efficient as directly querying the color class matrix for the corresponding row (i.e., constant time).

directions from parents to children in the tree starting from the dummy-node as the root for which we need a constant access from any node to all its adjacent ones. As we are storing the adjacency for a tree which is the most sparse possible graph representation for $n$ nodes, we would choose the adjacency list over matrix. Although the order for storing the tree adjacency list is $\mathcal{O}(n)$, the $n$ is considerably big and with high growth rate over samples that implementation-wise, the constant for the $n$ also matters. In a naive implementation, storing a weighted adjacency list of a tree with $n$ vertices assuming $k$ bytes to store an empty list in the language (at least 16 to store the start pointer and size) and the largest word size (8 bytes) to cover color-IDs greater than $2^{32}$ as well as a word size of 4 bytes for weights to cover number of samples greater than $2^{16}$ (both of which emerge in the scales we run Mantis on), requires $2 * n * (16 + 8 + 4)$ bytes. For example for indexing $80,000$ samples with $n \sim 4\text{e}9$, the required memory would be $\sim 208GB$. We note that this adjacency list should be in memory during the time filling the final structure of the MST which itself takes space.

We design a more thoughtful succinct representation of the adjacency list which eventually reduces the constant noticeably so that in practice the memory for that section is reduced in orders of magnitudes while still allowing constant-time access from each node to its neighbors. Assuming to have a tree with $n$ nodes where node IDs are in the range of $(0..n-1)$, we define the adjacency list through four succinct vectors; two of which are used to store information for the smaller end of an edge $\langle c_i, c_j \rangle$ and the other two for the larger end. We mention that since we do not have self-loops in the tree (edges with equal end IDs), therefore, for $n$ edges each of the four vectors are of size $n$. However, the width of the elements in each vector is different and basically the main reason that results in a total allocated space reduction.

In the first vector, $nei_{sm\_end}$, we store the IDs of the adjacent nodes of each node, if the node ID is smaller than its adjacent node ID, in sorted order of the node IDs along with the weight of the edge in a succinct form. Each word of the vector is of width $\log_2 n + \log_2 max(weights)$ bits where $max(weights) = s$, total number of samples. Since nodes can have different degrees as well as different number of connected nodes with the described condition, it is required to store the index of the start of the neighbor list for each node in $nei_{sm\_end}$. That takes us to the second vector, $start\_index_{sm\_end}$ in which at index $i$, we store the starting index of neighbor list for node $i$ in $nei_{sm\_end}$. In this way, to look up the neighbors of node $i$ and fetch the weights, we first fetch the start index of node $i$ at index $i$ of the $start\_index_{sm\_end}$, say its value is $start_i$; then jump to index $start_i$ in $nei_{sm\_end}$ for the first neighbor of node $i$. The count of the neighbors with greater ID value for each node $i$ is calculated by subtracting the start index of neighbors for $i$ and $i + 1$. If $start\_index_{sm\_end}[i] == start\_index_{sm\_end}[i + 1]$ this means $\nexists edge = \langle c_i, c_j \rangle \mid c_i < c_j$.

The other two vectors, $nei_{gr\_end}$, $start\_index_{gr\_end}$ follow the exact same results for storing the neighbors of node $i$ with IDs smaller than the ID of the node. In this way, we still are storing adjacency list for all the nodes in the tree to support constant-time access. However, we store the edge weights only in one of the vectors, for example in our case only in vector

$nei_{sm\_end}$. The memory consumption in this succinct representation would be $n*(\log_2 n*2+\log_2 max(weights))$ for the first pair vectors plus $n*(\log_2 n*2)$ for the second pair of vectors with total of $n*(4*\log_2 n+\log_2 s)$ which in the same example of indexing $80,000$ samples with $n \sim$ 4e9 would result in $\sim 67GB$. We can still improve this design by replacing the vectors indicating the start indices of neighbors for each node with a bit-vector of size $n$ and a rank data structure on top of it which in the same example would reduce the total memory down to $\sim 38GB$ which is $0.2^{th}$ of the naive implementation memory.

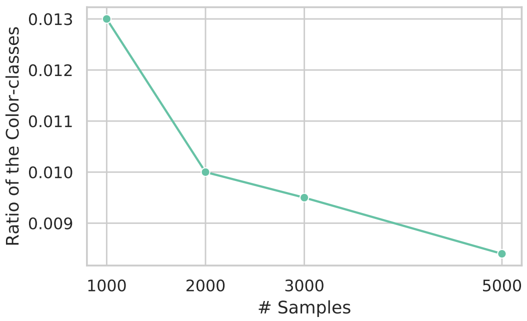


Figure 2: This plot shows the ratio of color-classes required to be stored in cache for merging two input indexes, based on number of RNA-sequencing samples that the output Mantis indexes when setting the max steps to walk the tree up to 16. As the plot shows, the ratio starts around 0.01 for an index on smaller number of samples which is way smaller than the max in theory ($\frac{1}{16}$). Even more interestingly, smaller number of color-classes than expected need to be loaded in cache to achieve the step threshold in query as we merging larger Mantis indexes.

### 3 Partitioned CQF Merge Pipeline

**Memory requirements:** Since the connection point of the left and right inputs are the minimizer blocks rather than the partitioned CQFs themselves, we do not need to load two partitioned CQFs into memory at the same time. We switch between inputs looking at the maximum minimizer covered by each input. If we have more minimizer blocks from left input, we load the next partitioned CQF from the right input and vice versa. This strategy guarantees that, at each point of the process, the total number of $k$-mers loaded into blocks does not pass twice the threshold for a partitioned CQF because at each point, we make sure that we can get rid of a subset of minimizer blocks by carefully choosing which input to process. Also, we would have at most two partitioned CQFs in memory when filling the output partitioned CQF (one input CQF and the output CQF). Altogether, the total memory requirement for the procedure based on the threshold $t$ for a partitioned CQF is $2 * t * sizeOf(kmer, colorID) + 2 * sizeOf(pcqf)$.

**Parallelization** : A single partitioned CQF is simply a CQF, and we can divide the range of hash values in a CQF into $t$ equal size segments ($t$ being number of threads given by the user) and let each thread separately walk the assigned segment and collect the $k$-mer and color pairs and partition them into their associated minimizer block. Walking the partitioned CQF and collecting the $k$-mers into different minimizers is performed many times throughout the merging process, and making this process work in parallel has a large effect in the final performance of the merge.

### 4 Commands and options used in running each of the experiments

In any of the commands that accepts multi-threading, the $< threadcount >$ option is set to "16".

#### 4.1 Mantis commands

Build Mantis index using iterative merge operation:

```
$ /usr/bin/time ./mantis build_by_merge -s 20 -t <
    thread_count> -p <process_count> -i <input_squeakr_list>
    -o <output_Mantis_index>
```

Merge two Mantis indexes:

```
$/usr/bin/time ./mantis merge -t <thread_count> -i1 <
    first_input_Mantis_index> -i2 <second_input_Mantis index>
     -o <output_Mantis_index>
```

Update Dynamic Mantis index:

```
$/usr/bin/time ./mantis lsmt_update -d <dynamic_Mantis_dir>
    -i <input_squeakr_list> -t <thread_count>
```

#### 4.2 VariMerge commands

Build Vari index (DBG and colors):

```
$/usr/bin/time ./cosmo-build -m <memory_limit> -d <
    input_kmc_list>

$/usr/bin/time ./pack-color <input_color_file> <
    number_of_colors> <TOTBITS> <SETBITS>
```

$< TOTBITS >$ and $< SETBITS >$ are derived from the logs of previous command, "cosmobuild"

Merge Vari indexes (VariMerge command)

```
$/usr/bin/time ./vari-merge <first_DBG> <second_DBG>

$/usr/bin/time ./color-merge merged.plan <
    first_COLOR_SD_VECTOR> <second_COLOR_SD_VECTOR> <
    first_number_of_colors> <second_number_of_colors>
```

#### 4.3 Bifrost commands

Build Bifrost over one/multiple raw sequencing sample(s) including all the $k$-mers:

```
$/usr/bin/time ./Bifrost build -r <input_file>/<
    input_file_list> -o <Bifrost_index_prefix> -t <
    thread_count> -c -k 23 -m 15
```

The input file can be of types Fastq or GFA. To discard any $k$-mer occurring only once in all files we replace "-r" with "-s"

We run Bifrost experiments in two modes, once we construct the final Bifrost directly over the list of Fastq files and once we construct a Bifrost index per each sample (which is a GFA file) and then construct the final Bifrost index over the GFA files. The first approach does not use any intermediate disk space but is relatively slower and requires more memory than the latter.

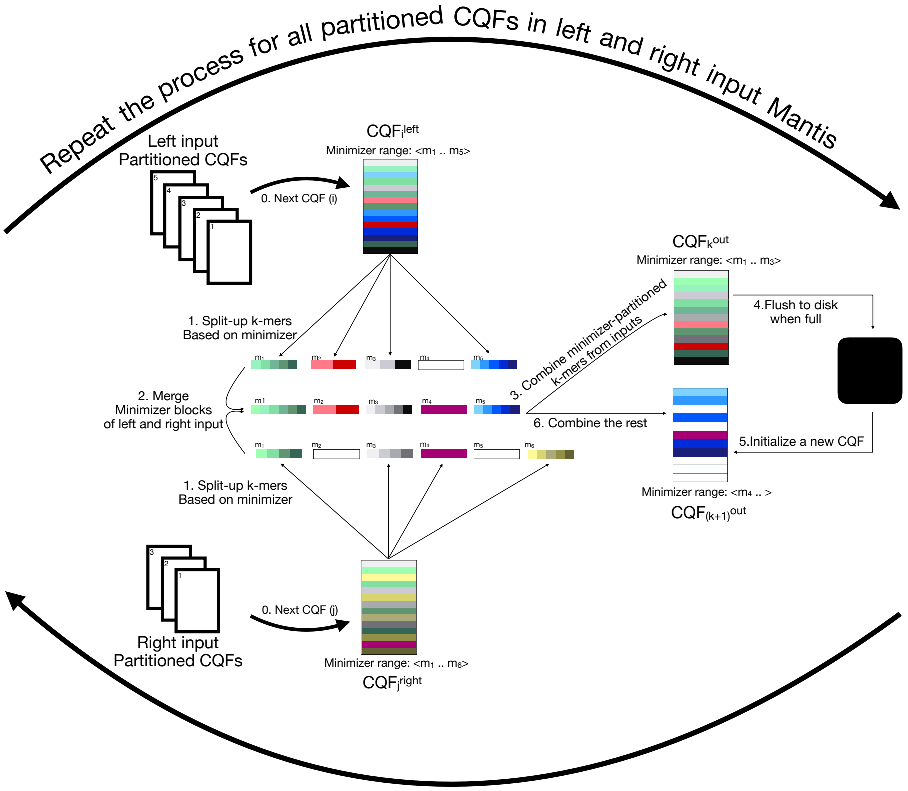


Figure 3: A toy example, illustrating the steps for merging two partitioned CQFs. The two arrows on top and bottom are indicative of the loop over each input partitioned CQF. For each partitioned CQF in each of the left and right inputs, first the $k$-mers are parted based on their minimizers. For each minimizer that its associated $k$-mers have been processed in both inputs, we merge the $k$-mers of the two input minimizer buckets. Then we walk over all the minimizer buckets for which we have the merged result, and insert the $\langle \mathrm{k}{-}mer, color \rangle$ pair into output CQF. Anytime the CQF is full, we flush it into disk, reinitialize a new partitioned CQF and continue inserting into the new partitioned CQF. We may reach the end of our merged minimizer buckets in the current round while our output CQF is not yet full; in that case, we continue filling it in next iterations.

**Algorithm 1** Merging two sets of partitioned CQFs from left and right CDBGs ($cdbg^l$ and $cdbg^r$) into output CDBG ($cdbg^o$)

```
colorMap : MAP(⟨color^out; ⟨color^left, color^right⟩⟩)
procedure MERGE_PCQFS(cdbg^l, cdbg^r, cdbg^o)
    COMPARE_SORTED_LIST(cdbg^l, cdbg^r, find_uniq_colorPairs)    ▷ Fills colorMap
    COMPARE_SORTED_LIST(cdbg^l, cdbg^r, store2cdbg)    ▷ Uses colorMap
end procedure

kcList^i_m: List of ⟨kmer, color⟩ pairs in a CQF partition for input cdbg^i where mnmzr(kmer) = m.
procedure COMPARE_SORTED_LIST(cdbg^l, cdbg^r, process)
    minMinimizer ← 0
    for p in MAX(# of CQF partitions for cdbg^l and cdbg^r) do
        for i in l, r do maxMinimizer^i ← cdbg^i.WALK_PARTITION(p, kcList^i)
        end for
        maxMinimizer ← MIN(maxMinimizer^l, maxMinimizer^r)
        for m in minMinimizer..maxMinimizer do
            kc^l = kcList^l_m.GETCURRENT()    ▷ kc = ⟨kmer, color⟩
            kc^r = kcList^r_l.GETCURRENT()

            repeat
                if kc^l.kmer < kc^r.kmer then
                    process(kc^l, NA)    ▷ NA: Not Available
                    kc^l.NEXT()
                else if kc^l.kmer > kc^r.kmer then
                    process(NA, kc^r)
                    kc^r.NEXT()
                else
                    process(kc^l, kc^r)
                    kc^l.NEXT()
                    kc^r.NEXT()
                end if
            until kc^l.HASNEXT() or kc^r.HASNEXT()

            DELETE(kcList^l_m)
            DELETE(kcList^r_m)
        end for
        minMinimizer ← maxMinimizer
    end for
end procedure
```

```
procedure FIND_UNIQ_COLORPAIRS(kc^l, kc^r)                ▷ kc = ⟨kmer, color⟩
    if ⟨kc^l.color, kc^r.color⟩ ∉colorMap then
        index ←colorMap.length
        colorMap.ADD(⟨kc^l.color, kc^r.color⟩ → index )
    end if
end procedure
procedure STORE2CDBG(kc^l, kc^r)                          ▷ kc = ⟨kmer, color⟩
    colorID ←colorMap [⟨kc^l.color, kc^r.color⟩]
    if kc^l! = NA then
        kmer ← kc^l.kmer
    else
        kmer ← kc^r.kmer
    end if
    cqf^o ← cdbg^o.CURRENTPARTITION()
    if cqf^o.ISFULL then
        cqf^o.STORE2DISK()
        cqf^o ← cdbg^o.NEWPARTITION()
    end if
    cqf^o.INSERT(⟨kmer, color⟩)
end procedure
l : minimizer length, “8” in our case
procedure WALK_PARTITION(p, kcList)                       ▷ p: partition ID
    pcqf_p ← LOAD p^th CQF partition from disk
    if pcqf_p == NULL then
        return MAXINT
    end if
    for kc in pcqf_p do
        m ← FIND_MINIMIZER(kc.kmer)                       ▷ (Between 0 and 4^l)
        kcList_m.INSERT(kc)
        maxMinimizer ←MAX(maxMinimizer, m)
    end for
    return maxMinimizer
end procedure
```
